# A thermoreversible hydrogel model for *in situ* and *ex situ* analysis of tumor-immune interactions

**DOI:** 10.64898/2026.11.15.766807

**Authors:** Carmen Escalona-Noguero, Nina Anseeuw, Guillermo Solís-Fernández, Steven Huysecom, Sandra P. Nunes, Behrad Shaghaghi, Paul Kouwer, Ilaria Elia, Susana Rocha

## Abstract

Despite the clinical success of cancer immunotherapy, many patients fail to respond or develop resistance, highlighting the need to understand how interactions between tumor and immune cells influence treatment outcomes. Three-dimensional in vitro models allow these interactions to be studied under controlled conditions. However, Matrigel and other biologically derived scaffolds can vary in composition, offer limited independent control over matrix properties, and require cell-recovery procedures that complicate downstream analysis. Here, we present a co-culture model of melanoma spheroids and primary CD8^+^ T cells based on a defined, thermoreversible synthetic polyisocyanopeptide (PIC) hydrogel. PIC supported T cell viability and migration, with mean motility parameters similar to those in Matrigel and lower variability between experiments. Fluorescence imaging revealed CD8^+^ T cell infiltration and increased melanoma-cell apoptosis during co-culture. Cooling enabled recovery of nearly all embedded cells in less than 3 min without chemical or enzymatic matrix digestion. Metabolic profiling of recovered T cells identified time-dependent changes in intracellular lactate, amino acids, and tricarboxylic acid cycle intermediates, alongside changes in extracellular metabolite availability. Flow cytometry distinguished T cell populations with different effector profiles, with higher expression of effector markers in the fraction associated with melanoma spheroids. By combining a defined synthetic matrix with *in situ* imaging and rapid recovery for *ex situ* analysis, this model connects the spatial dynamics of tumor-immune interactions with metabolic and immune phenotypes, providing a basis for studies of immune-cell function and responses to immunotherapy.

## INTRODUCTION

Immunotherapy has changed the treatment landscape of several types of cancer, including melanoma, lung cancer, and hematologic malignancies[1–4]. Whereas chemotherapy and radiotherapy act primarily by killing cancer cells or inhibiting their growth, immunotherapy harnesses the immune system’s intrinsic ability to eliminate them[5,6]. In advanced melanoma, for instance, immunotherapy has become a standard treatment, with combined immunotherapy achieving a 10-year overall survival rate of 43% in patients with unresectable or metastatic disease[7]. Many current immunotherapies act by enhancing the activity of cytotoxic CD8^+^ T cells, which are effector cells of the immune system able to recognize and kill malignant cells. Despite the reported success, many patients fail to respond to treatment or subsequently develop resistance, and the mechanisms determining treatment success or failure remain incompletely understood[8–10]. Several studies have identified CD8^+^ T cell exhaustion and metabolic constraints within the tumor microenvironment (TME) as contributors to impaired antitumor activity[11]. Low oxygen availability has been linked to T cell dysfunction in mouse melanoma models, while tumor-derived lactate can alter T cell metabolism and reduce cytotoxicity[12], [13], [14]. Understanding how the TME suppresses T cell responses requires models in which tumor-immune interactions can be manipulated and followed over time. *In vivo* models provide a physiological context, but they often do not allow tumor-immune interactions to be dissected in the depth required for mechanistic characterization. Moreover, it is difficult to conduct individual manipulations without affecting other aspects of the TME. In contrast, *in vitro* systems offer greater control over cell composition and biochemical conditions, together with direct access to readouts such as T cell migration, tumor infiltration, cancer-cell killing, and immune-cell phenotype[15]. However, mimicking the properties of the TME remains challenging.

In this context, 3D co-culture models combining tumor spheroids and immune cells recapitulate key features of the TME that cannot be captured by 2D co-cultures, such as nutrient and oxygen gradients and spatially organized tumor-immune interactions. For example, collagen-embedded melanoma spheroids enabled live measurements of recruitment, infiltration, and killing by cytokine-induced killer cells and NK-92 cells[16]. Microfluidic cultures of melanoma spheroids in Matrigel further revealed that cytotoxic T cells accumulated at sites of spheroid fragmentation, providing evidence that cooperation between T cells facilitates tumor destruction[17]. Beyond measuring cancer-cell death, spheroid co-cultures can also reveal changes in the immune cells themselves. After culturing antigen-presenting melanoma spheroids in Matrigel with primary human cytotoxic T cells, recovered T cells showed reduced killing capacity and interferon-γ secretion, demonstrating that tumor contact can induce functional suppression *in vitro*[18]. These approaches link the spatial dynamics of tumor-immune interactions to cellular responses, but both cell behavior and the range of possible readouts depend strongly on the properties of the surrounding scaffold[19], [20], [21].

The scaffold, or surrounding matrix, provides the environment through which immune cells must migrate before reaching the tumor spheroid, making its physical and biochemical properties relevant to the interactions being studied[22]. Matrigel and other naturally derived matrices are widely used for 3D cultures, although batch-to-batch variability and limited control over individual matrix properties complicate their use in mechanistic studies[22], [23], [24]. Changing matrix concentration, for example, alters both pore structure and adhesive ligand density, often alongside stiffness, making their individual effects on cell behavior difficult to distinguish. Synthetic hydrogels offer more defined compositions and allow biochemical and mechanical properties to be controlled more independently[25]. However, many established synthetic systems form covalently crosslinked networks. Depending on their pore size and degradability, these networks can restrict cell movement by limiting the space available for migration and the ability of cells to locally displace or remodel the matrix. Furthermore, for *ex situ* analysis, recovering cells from covalently crosslinked hydrogels often requires chemical or enzymatic degradation of the surrounding matrix. Prolonged processing or exposure to digestion reagents can affect cell viability, surface markers, and functional state, while delays in recovery can alter the metabolic state being measured. Supramolecular hydrogels offer an alternative in which non-covalent interactions confer dynamic properties on the network[26]. Depending on the strength and lifetime of these interactions, local network rearrangement can accommodate cell movement without requiring irreversible matrix degradation.

A particularly notable example of supramolecular hydrogels used in 3D culture are the polyisocyanopeptide (PIC) gels, an established synthetic material in which polymer chains assemble into a fibrous network through non-covalent interactions[25], [27], [28]. Their defined composition and tunable mechanical properties allow the scaffold to be tuned, avoiding complex mixtures of biological components and growth factors present in Matrigel[29]. The fibrous architecture of PICs gels provides an extracellular matrix-like environment that supports cell migration and interactions. Moreover, cell behavior can be easily monitored over time due to the optical transparency of the material. Especially relevant to the application here presented is the thermoreversible nature of PIC: it forms a hydrogel upon warming and returns to a liquid upon cooling. This allows rapid cell recovery by adding ice-cold buffer followed by centrifugation, without chemical or enzymatic degradation of the matrix, which is key for certain downstream analysis, such as metabolic profiling.

Here, we used the optical transparency and thermoreversibility of PIC hydrogels to develop a three-dimensional (3D) co-culture model of melanoma spheroids and primary CD8^+^ T cells, combining *in situ* imaging with *ex situ* analysis of recovered cells. To study these interactions in melanoma, a cancer widely treated with immunotherapy, we generated spheroids from the murine cell lines M10M5 OVA and M171M1 OVA, which carry BRAF and NRAS mutations, respectively[30]. These mutations define two major genetic subtypes of melanoma, allowing us to assess T cell infiltration and tumor-cell killing across distinct melanoma backgrounds. Both cell lines express ovalbumin (OVA) as a model antigen and were co-cultured with primary CD8^+^ T cells isolated from OT-I mice. These T cells recognize the OVA-derived peptide SIINFEKL presented by major histocompatibility complex class I (MHC-I) molecules. Upon recognition, OT-I T cells can kill OVA-expressing melanoma cells, providing a defined system for following antigen-specific T cell interactions with tumor spheroids and assessing their cytotoxic activity.

Using this model, we monitored CD8^+^ T cell migration, spheroid infiltration, and melanoma cell death by fluorescence imaging, and then recovered the T cells to analyze their polar metabolites by mass spectrometry and their immune phenotype by flow cytometry. By combining a defined synthetic matrix with *in situ* imaging and straightforward access to recovered cells, this approach connects the spatial dynamics of tumor-immune interactions with changes in immune-cell phenotype. The model here developed provides a basis for future studies of T cell dysfunction, metabolic constraints, and responses to immunotherapy in melanoma, with potential for adaptation to other solid tumors and immune-cell populations.

## RESULTS

### PIC hydrogels support CD8^+^ T cell viability and migration

Before establishing co-cultures with melanoma spheroids, we assessed whether PIC hydrogels maintained CD8^+^ T cell viability and allowed migration through the surrounding matrix. We used PIC at a concentration of 1 mg/mL to obtain a stable gel with relatively large pores[26]. Throughout the manuscript, the abbreviation PIC refers to this formulation unless otherwise stated. We examined cell viability following recovery, cellular morphology, and cell motility within PIC hydrogels, using suspension culture and Matrigel as reference conditions.

CD8^+^ T cells were cultured in PIC for 24 h, recovered, and assessed for viability using Trypan blue exclusion. Cell viability was comparable to that of cells maintained in suspension, with no significant difference between the two conditions (Fig. 1A). This was further confirmed by flow-cytometry analysis using a viability dye (Fig. 1A). We next examined cell morphology after 24 h in PIC or Matrigel. Cells were fixed and stained to visualize nuclei and filamentous actin, and morphology was quantified using cell size, aspect ratio, and solidity (Fig. 1B). For the different parameters, there was no significant difference between T cells cultured in PIC or Matrigel. Cells were slightly larger when grown in PIC, but the difference was not significant.

**Figure 1:**
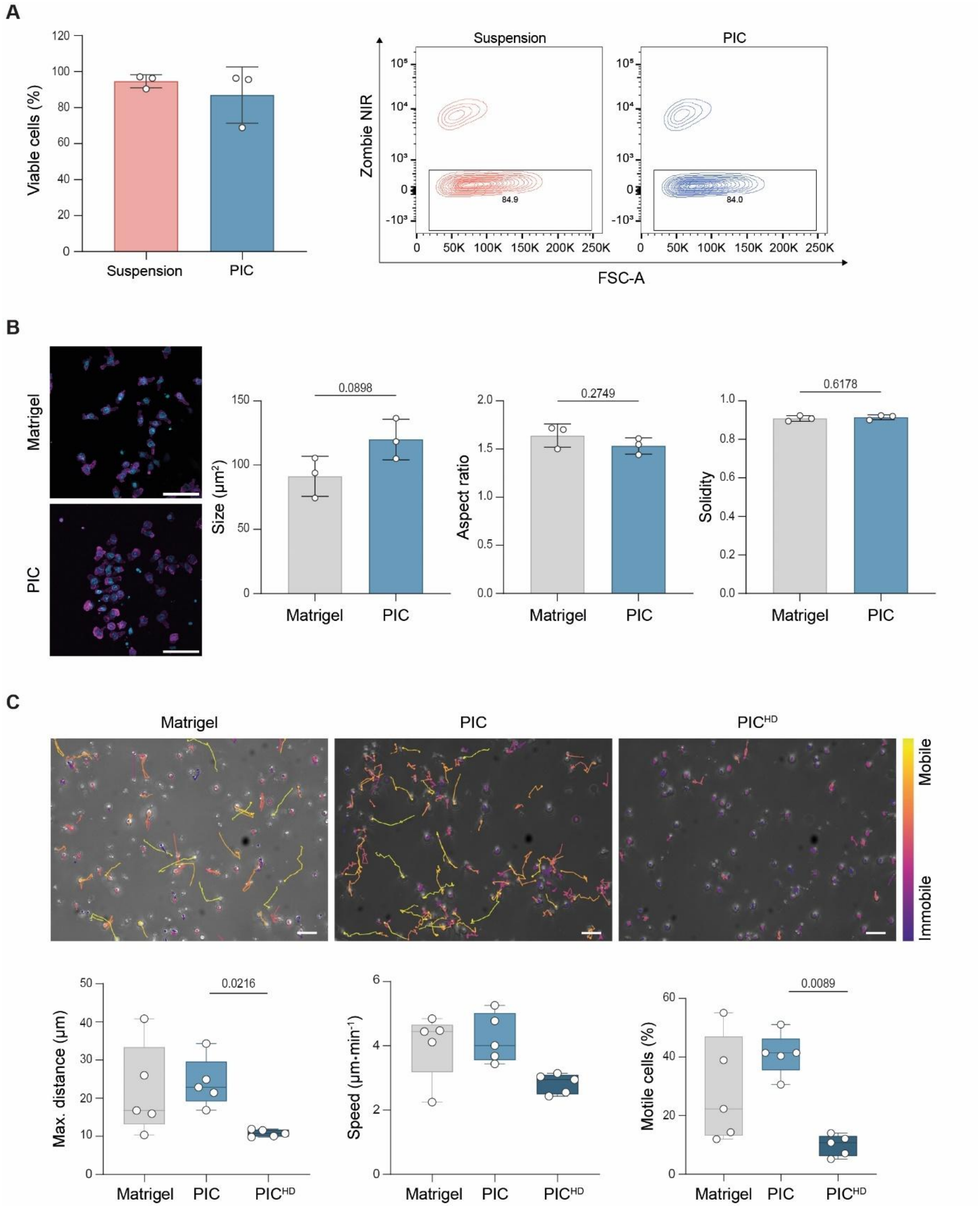
CD8^+^ T cell viability, motility, and morphology in PIC. (a) Viability after 24 h in PIC was assessed by trypan blue exclusion and automated counting (left) and by Zombie NIR live/dead staining and flow cytometry (right). Cells cultured in suspension for 24 h served as a control. (b) Characterization of CD8^+^ T cell morphology in PIC. Nuclei (Hoescht, blue) and actin cytoskeleton (phalloidin, magenta) were stained to determine cell area, aspect ratio, and solidity. Cells embedded in Matrigel were used as control. n=3 biological replicates. Scale bar: 50 μm. (c) Representative images and quantification of CD8^+^ T cell motility in Matrigel, PIC and PIC^HD^ (high density) assessed by live brightfield imaging. The color of the tracks represents the degree of motility of the cell. n=3 biological replicates. Scale bar: 50 μm

To determine whether T cells could migrate through PIC, we tracked their movement in 1 mg/mL PIC and Matrigel. We also included 2 mg/mL PIC (high density, PIC^HD^) to examine how increasing network density affected motility (Fig. 1C). This increase in concentration produces a small change in mechanical properties and a larger reduction in pore size[26]. The mean fraction of mobile cells was similar in PIC and Matrigel, although Matrigel showed greater variability between independent experiments (Fig. 1C). This variability is relevant when comparing T cell behavior across experimental conditions, as it can make smaller biological effects harder to distinguish. Variations in composition and mechanical properties within and between Matrigel batches have previously been reported and may contribute to this inconsistency[31]. In contrast to both PIC and Matrigel, T cells in PIC^HD^ showed reduced diffusion and a smaller fraction of mobile cells (Fig. 1C). This concentration dependence is consistent with increased physical confinement in the denser network, although a contribution from differences in mechanical properties cannot be ruled out.

Together, these results show that PIC maintains CD8^+^ T cell viability over 24 h and permits cell migration, supporting its use for subsequent co-culture experiments.

### PIC-based co-culture models enable *in situ* analysis of T cell infiltration and tumor-cell death

Having established that PIC supports CD8^+^ T cell viability and migration, we next examined whether this matrix allows T cells to interact with melanoma spheroids, infiltrate their multicellular structure, and induce tumor-cell apoptosis. We first characterized spheroid size and morphology to provide a reference for interpreting the subsequent co-culture experiments.

Spheroids were generated from M10M5 OVA (*Braf*^*V600E*^; *Pten*^*Δ/Δ*^) and M171M1 OVA (*Nras*^*Q61R*^; *Pten*^*Δ/Δ*^) murine melanoma cells (Fig. 2A) and characterized by measuring their volume (Fig. 2B). We then embedded spheroids in PIC, either alone or together with primary OT-I CD8^+^ T cells and followed cell behavior by time-lapse bright-field microscopy. In the absence of T cells, melanoma cells invaded the surrounding PIC, forming protrusions that extended from the spheroid (Fig. 2C; Supplementary Movies 1-2). In contrast, these invasive protrusions were not observed in co-culture conditions, and spheroids maintained or decreased their size over the observation period (Fig. 2C; Supplementary Movie 3-4). The absence of invasive protrusions in co-culture is consistent with T cells targeting melanoma cells as they emerge from the spheroid, although killing of these individual cells could not be established from bright-field imaging alone. To measure T cell killing activity in the co-culture, we quantified cell death using PI staining. 30 h after T cell addition, the area ratio of PI fluorescence within the spheroid was significantly higher in co-cultured spheroids than in spheroids cultured without T cells (Fig. 2D). This result is consistent with increased melanoma-cell apoptosis. Because bright-field images do not reliably distinguish T cells from melanoma cells, we also used CD8 immunostaining and confocal microscopy to identify the T cells and determine whether they were able to infiltrate the spheroids. As shown in Fig. 2E, CD8^+^ T cells were detected within the melanoma spheroids after 36 h of co-culture, indicating that T cells can infiltrate the spheroids.

**Figure 2:**
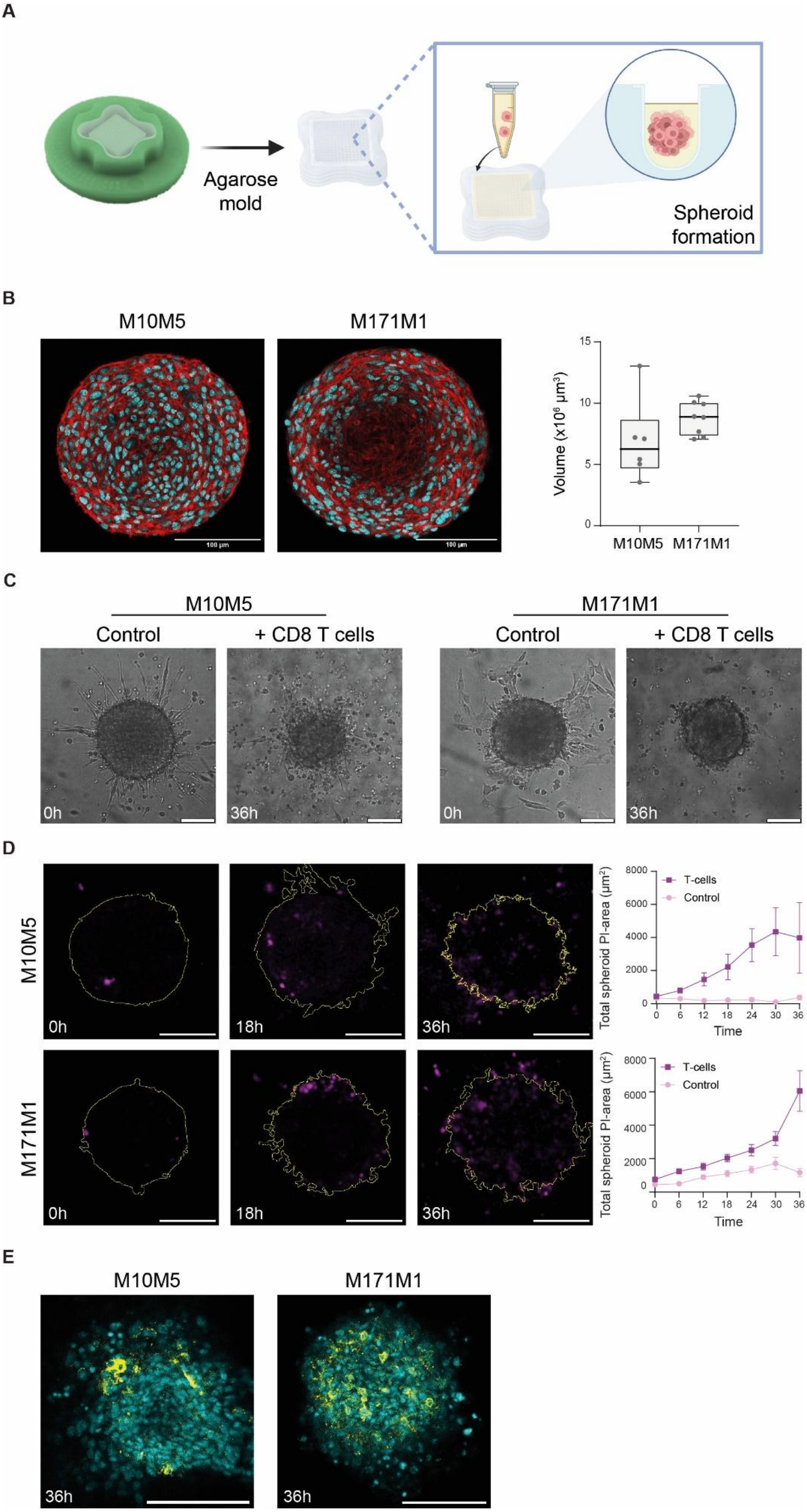
co-culture of CD8^+^ T cells and melanoma spheroids in PIC and functional in situ analysis. (a) Schematic of the procedure followed to generate M10M5 and M171M1 spheroids using agarose micromolds. (b) Spheroid structure and volume analysis. Hoescht-stained nuclei are shown in cyan and the actin cytoskeleton, stained with phalloidin, in red. Scale bar: 100 μm. (c) Representative end-point images (36 h) of M10M5-OVA and M171M1-OVA spheroids embedded in PIC, alone or in co-xculture with OT-I CD8^+^ T cells. Scale bar: 100 μm. (d) Representative images and time-course quantification of T cell-mediated melanoma cell killing in PIC co-cultures. PI-positive cells are shown in magenta. n=1 biological replicate. Scale bar: 100 μm. (e) Infiltration of CD8^+^ spheroids (yellow) in the spheroid structure after 36 h of co-culture in PIC. Hoescht-stained nuclei are shown in cyan. Scale bar: 100 μm.

Together, these experiments establish the PIC-based co-culture as a model for following tumor-immune interactions through complementary imaging readouts. The matrix enables live imaging of melanoma cell invasion, immunostaining-based localization of infiltrating T cells, and quantification of tumor cell killing, allowing immune and melanoma cell behavior to be monitored within a single 3D culture system.

### Rapid recovery from PIC enables *ex situ* metabolic analysis and immune profiling of T cells

In addition to its biocompatibility and optical transparency, PIC offers the advantage of thermoreversible gelation, enabling rapid T cell recovery without chemical or enzymatic matrix digestion. To demonstrate how this property extends the model’s analytical capabilities beyond imaging, we recovered T cells after co-culture for metabolic analysis and flow-cytometry profiling of their immune phenotype. For *ex situ* analysis, co-cultured samples were cooled by adding ice-cold PBS, and cells were collected by differential centrifugation. This procedure recovered nearly 100% of the embedded cells in less than 3 min, minimizing processing time before metabolic analysis (Fig. S1).

We next examined changes in the metabolic profile of CD8^+^ T cells during co-culture with M10M5 melanoma spheroids (Fig. 3A). T cells were collected before embedding them in the PIC matrix (0 h) and after 12, 24, and 36 h of co-culture; and their polar metabolites were extracted and analyzed by gas chromatography mass spectrometry (GC-MS). (Fig 3A). We observed that the co-culture induced marked and often time-dependent changes in glycolysis-related metabolites, amino acids, and TCA intermediates. Intracellular lactate, the main by-product of glycolysis, increased progressively over the 36 h, reaching approximately 16-fold the level measure before co-culture. In turn, the levels of pyruvate, the end-product of glycolysis, peaked at 12 h and decreased after that. Together, these changes could indicate increased glycolytic activity. Alanine and lysine also accumulated steadily over time, whereas serine and proline increased markedly at 12h and decreased slightly at later time points. Other amino acids such as glycine, aspartate, and glutamate showed minimal variations, with only a modest decrease in glutamate. The levels of TCA metabolites (alpha-ketoglutarate (a-KG), fumarate and malate) decreased between 12 and 36 h. Overall, these data suggest an early metabolic shift in T cells after co-culture onset, with accumulation of glycolytic products, accompanied by changes amino acids and TCA cycle intermediates.

**Figure 3:**
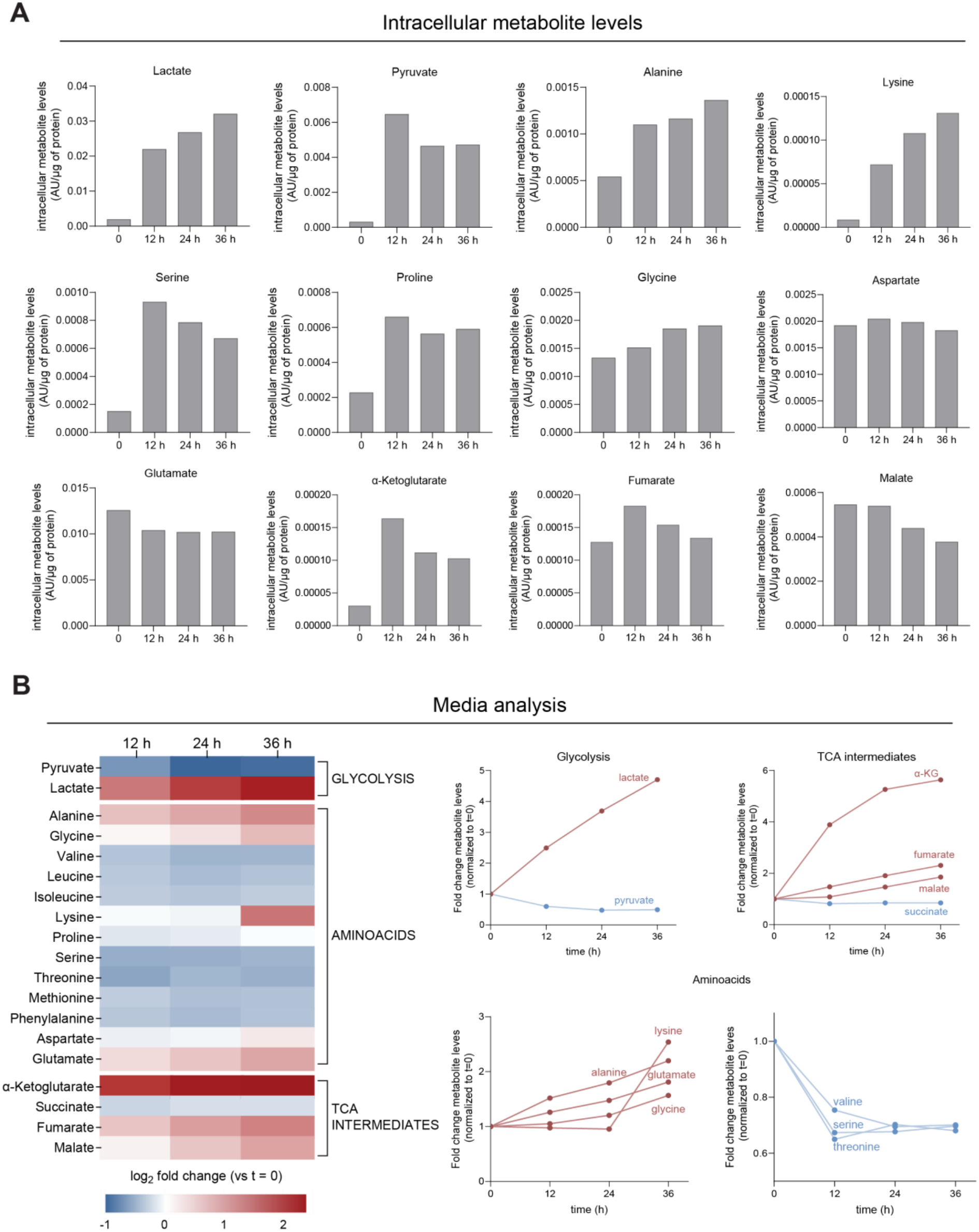
Time-course metabolic profiling of CD8^+^ T cells recovered from co-culture and their extracellular medium. (a) Relative levels of intracellular polar metabolites in OT-I CD8+ T cells co-embedded with M10M5-OVA spheroids in PIC. n=1 biological replicate. (b) Relative metabolite concentrations in the extracellular media normalized to t=0. n=1 biological replicate.

To complement the measurements of intracellular metabolite levels, we quantified changes in the composition of the extracellular media (Fig. 3B). Again, we observed variations in glycolysis-associated metabolites, amino acids, and TCA intermediates. Pyruvate was gradually depleted from the media, whereas lactate accumulated progressively over the 36-hour period, with its concentration increasing nearly 5-fold. The accumulation of lactate, as previously mentioned, is one of the metabolic hallmarks of the tumor microenvironment due to the high glycolytic rate of cancer cells. In addition, in line with the existing literature, we observed changes in amino acid availability[32], [33]. Glutamate, alanine, glycine, and lysine concentrations were enriched over time, while serine, valine, and threonine were depleted in the first 12 hours of co-culture and remained stable afterwards. Finally, we also observed an accumulation of the TCA cycle metabolites fumarate, malate, and, especially, alpha-ketoglutarate.

Finally, we used flow cytometry to examine whether T-cell phenotype differed according to their association with melanoma spheroids. Following PIC liquefaction, differential centrifugation yielded two fractions: T cells that co-sedimented with the spheroids, termed proximal CD8^+^ T cells, and those remaining in suspension, termed distal CD8^+^ T cells (Fig. 4A). These designations describe the fractions obtained during recovery rather than the cells’ precise locations during co-culture. The expression levels of effector markers were analyzed by flow cytometry (Fig. 4B). After 36 h of co-culture, we observed that spheroid-proximal T cells showed a higher expression of interleukin 2 (IL-2), tumor-necrosis factor α (TNFα), and granzyme B (GzmB) compared to distal T cells, suggesting that the proximity to the spheroid induces a more effector-like profile. These differences were more marked in the case of M10M5, suggesting that the different oncogenic mutations of these cell lines might have an impact on T cell activity.

**Figure 4:**
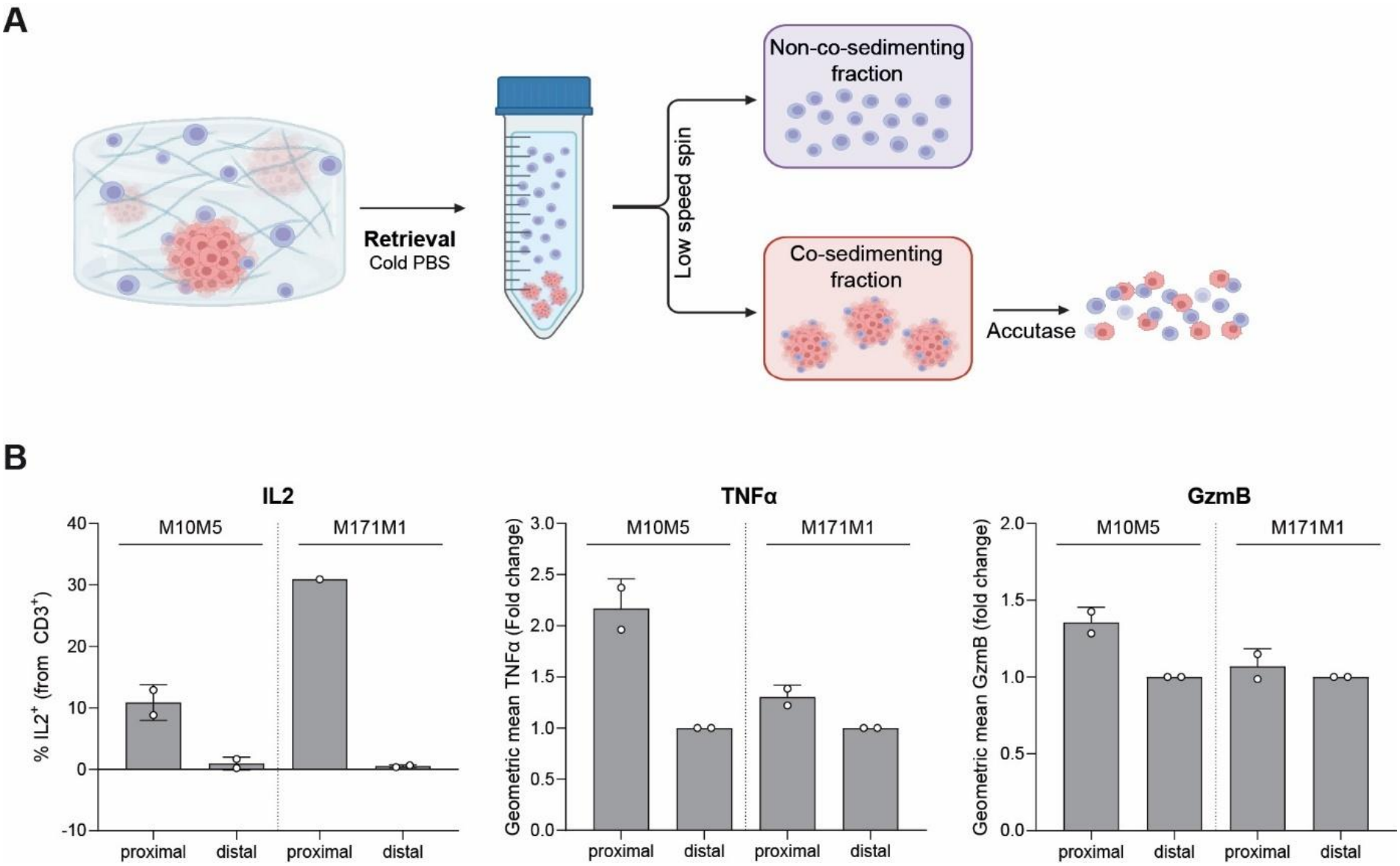
Immune profiling of spheroid-proximal and spheroid-distal CD8^+^ T cells recovered from co-culture. (a) Schematic of the procedure followed to recover spheroid-proximal and distal CD8^+^ T cells from the PIC co-culture with M10M5-OVA and M171M-OVA spheroids. (b) Flow cytometry analysis of effector markers in the two T cell populations. n=2 biological replicates.

Together, these findings demonstrate that rapid recovery from PIC enables metabolic and immune profiling of CD8^+^ T cells after three-dimensional co-culture, alongside measurements of the extracellular metabolic environment. The defined, tunable matrix connects *in situ* imaging of tumor-immune interactions with *ex situ* analysis of cellular responses, providing a basis for examining how culture conditions affect T-cell metabolism and effector phenotype.

## CONCLUSIONS

We established a three-dimensional co-culture model in which a synthetic PIC hydrogel supports primary CD8^+^ T cell migration, melanoma spheroid infiltration, and imaging of tumor-cell apoptosis. T cell motility was similar to that measured in Matrigel, with lower variability between experiments, while the thermoreversible network enabled rapid cell recovery without chemical or enzymatic matrix digestion. This combination allowed *in situ* observations to be complemented by *ex situ* analysis of T cell metabolic and immune profiles. By combining a defined synthetic matrix with live imaging and rapid cell recovery, the model connects the spatial dynamics of tumor-immune interactions with cellular responses while limiting the uncontrolled biochemical cues introduced by complex, biologically derived scaffolds.

From a biomaterials perspective, these findings highlight the importance of considering matrix properties and cell recovery together when designing three-dimensional culture systems. A scaffold must accommodate cell movement and interactions while remaining compatible with the measurements needed to interpret them. PIC offers opportunities to vary mechanical properties and adhesive cues in future studies, although changes in network architecture and their effects on cell access will require characterization. The model could also be extended to other solid tumors, stromal populations, and immune cells to examine how the local environment affects antitumor activity. Such developments would support studies of immune-cell dysfunction and immunotherapy responses, building on the migration, infiltration, imaging, and recovery capabilities established here.

## MATERIALS AND METHODS

### CD8^+^ T cell isolation and activation

Primary naïve CD8^+^ T cells were isolated from the spleens of wild-type C57BL/6J and OT-I mice using Biolegend’s naïve CD8^+^ T cell isolation kit (480035), according to the instructions of the manufacturer. Naive CD8^+^ T cells were then activated using plates coated with 2 μg/mL of anti-CD3 (BioXCell, BE0001-1, clone 145-2C11) and anti-CD28 (BioXCell, BE0015-1, clone 37.51). Cell density at seeding was 1-1.5 million cells/mL. T cells were activated during 48 hours in RPMI 1640 (Gibco, 21875034) supplemented with 10% foetal bovine serum (Biowest, S181B), 100 U/mL penicillin, 100 mg/mL streptomycin (Gibco, 11528876), 10 mM HEPES (Gibco, 12509079), 1 mM sodium pyruvate (Gibco, 12539059), 55 mM beta-mercaptoethanol (Gibco, 11508916),100 U/mL IL-2 (R&D Systems, 202-IL-050) and 10 ng/mL IL-12 (PeproTech, 210-12-50 mg).

### Cancer cell lines

M10M5 and M171M1 murine melanoma cell lines were generated by the laboratory of Florian Karreth (H. Lee Moffitt Cancer Center)[30]. OVA-expressing M10M5 and M171M1 were generated by viral transduction using a lentiviral vector with the open reading frame under the control of the CMV (cytomegalovirus) promoter. Cells were grown in RPMI 1640 (Gibco, 21875034) supplemented with 10% foetal bovine serum (Biowest, S181B), 100 U/mL penicillin, 100 mg/mL streptomycin (Gibco, 11528876), 10 mM HEPES (Gibco, 12509079), 1 mM sodium pyruvate (Gibco, 12539059), 55 mM beta-mercaptoethanol (Gibco, 11508916). All cell lines were confirmed Mycoplasma negative through monthly PCR-based tests.

### PIC polymers

RGD-functionalized 1K PIC hydrogels were prepared as previously described by Yuan *et al*[34]. Briefly, azide-functionalized PIC was functionalized with DBCO-PEG4-GRGDS using click chemistry. The functionalized polymer was purified by precipitation, dried and stored at -20 °C until use. PIC hydrogels were prepared by dissolving the polymer in PBS at 4 °C overnight before use, respectively PIC at 2 mg/mL and PIC^HD^ at 4 mg/mL.

### Cell viability assay

T cells were embedded in PIC hydrogels and incubated at 37 °C for X minutes. Cells were then recovered from the PIC by adding cold PBS and their viability was determined by trypan blue exclusion using an automated cell counter. Cells in suspension (non-embedded) were used as viability control. Results were confirmed by flow cytometry upon staining with Zombie NIR (BioLegend, 423105, 1:4000 dilution). Further details can be found in the *Flow cytometry* section.

### Motility assay

T cells were embedded in Matrigel (7:10 hydrogel-cell ratio) or PIC hydrogels (1:1 ratio) at a final concentration of 1 million cells/mL. PIC hydrogels were used at final polymer concentrations of 1 mg/mL (PIC) or 2 mg/mL (PIC^HD^). T cell motility was imaged for 30 min using the bright field channel on a Leica SP8 confocal microscope equipped with an environmental control system (The Cube chamber heater and The Brick CO_2_ gas mixer (Life imaging Services)). T cells were maintained in motility medium consisting of RPMI supplemented with glucose and glutamine, BSA and HEPES, without exogenous interleukins, as described in Simula *et al*[35]. For the analysis, cell segmentation was performed in Napari using a trained segmentation model, followed by automated tracking using Trackastra. Tracks were subsequently analyzed in MATLAB using mean squared displacement (MSD) to quantify and classify T cell motility. The anomalous diffusion exponent (α) was used to classify cell motion as active (α > 1) or subdiffusive/confined (α < 1). Tracks containing five or fewer data points were excluded from the analysis.

### Analysis of CD8^+^ T cell size and morphology

CD8^+^ T cells were embedded in Matrigel and PIC following the same procedure as for the motility assay. 24 hours later, they were fixed with 4% paraformaldehyde (PFA) and stained with Hoescht and phalloidin-647 to visualize their nuclei and actin cytoskeleton, respectively. Cell pose was used to segment cells and analyze their size, aspect ratio (ratio between the long and short axes) and solidity (shape regularity).

### Co-culture of OT-I CD8^+^ T cells and M10M5/M171M1-OVA spheroids in PIC

CD8^+^ T cells and spheroids were co-embedded in PIC hydrogels at a final T cell concentration of 2 million cells/mL, using a volume ratio of 2:1:1 (PIC:spheroids:CD8+ T cells). The samples were incubated at 37 °C to allow hydrogel gelation.

### Killing assay

Following gel formation, pre-warmed propidium iodide (PI) staining solution (1:7500 dilution in RPMI supplemented with IL-2 and IL-12) was added on top of the hydrogel. In situ imaging of the co-culture was subsequently performed using a Leica TCS SP8 confocal microscope equipped with an environmental control system to maintain physiological conditions (37 °C and controlled CO_2_ levels). PI and bright-field (BF) signals were acquired at hourly intervals over a 36-hour period to monitor spheroid killing by CD8^+^ T cells.

### Evaluation of CD8^+^ T infiltration in melanoma spheroids

After 36 hours in co-culture in PIC cells were fixed with 4% PFA for 1 hour at 37 °C and then permeabilized and stained following the procedure described in Yuan *et al*[34]. CD8^+^ cells were stained with anti-mouse CD8a primary antibody (Proteintech, 65205-1-Ig, dilution 1:2000) and Goat anti-Rat IgG Alexa Fluor™ 647 secondary antibody (ThermoFisher Scientific, A-21247). Nuclei were stained with Hoechtst.

### Harvesting of CD8^+^ T cells from the 3D co-culture for FACS analysis

T cells and spheroids were recovered from the PIC by adding 100 uL of cold PBS to each well (96-well plate). Per replicate, 4-6 wells were pooled together to ensure that the number of cells was enough for downstream analysis. To separate spheroid-proximal and spheroid-distal CD8^+^ T cells, the mix was subjected to a short low speed spin (∼10 seconds) and the supernatant (distal T cells) was transferred to a separate tube and washed once with cold PBS. The pellet, containing the spheroids and the T cells that co-sediment with them (proximal T cells) was washed once with PBS and then subjected to enzymatic dissociation with Accutase (Thermofisher, 00-4555-56) for 5-10 minutes at 37 °C.

### Flow cytometry

To enable cytokine detection, after 36 h in co-culture cells were stimulated with 100 ng/mL phorbol 12-myristate 13-acetate (Sigma, P1585-1MG) and 500 ng/mL ionomycin (Sigma, I0634-1MG) in the presence of GolgiStop (BD Biosciences, 554724) for 3 hours at 37 °C. Cells were then harvested from the 3D co-culture as described above, washed with PBS and stained with Zombie NIR Fixable Viability Kit (BioLegend,. 423105, 1:4000 dilution) for 20 minutes at room temperature to exclude dead cells. Next, they were resuspended in staining buffer (PBS supplemented with 1 % FBS and 2 mM EDTA) and stained with the indicated antibodies for 30 minutes at 4 °C. Antibodies: BV480 anti-mouse CD3 (eBioscience, 414-0032-80 ); R718 anti-mouse CD8b (BD, 566985 ); PE-Dazzle 594 anti-mouse CD45 (BioLegend, 103145 ); BV711 anti-mouse PD-1 (BioLegend, 135231); APC anti-mouse TIM-3 (eBioscience, 17-5870-82);

BV650 anti-mouse Slamf6 (BD, 740628); BV421 anti-mouse IFNg (BioLegend, 505829); BV785 anti-mouse TNFa (BioLegend, 506341); PE-Cy7 anti-mouse GzmB (Biolegend, 506341)., FITC anti-mouse IL-2 (Biolegend; 503805 ). Flow cytometry was performed on BD FACSymphony A1 Cell Analyzer (Waters Biosciences) and FlowjoTM 10.10.1 software (Waters Biosciences) was used for data analysis. UltraComp beads (Invitrogen; 01222242) were used for compensation.

### Harvesting of CD8^+^ T cells from the 3D co-culture for metabolite extraction

T cells and spheroids were recovered from the PIC by adding 100 uL of cold PBS to each well (96-well plate). Per replicate, 6 wells were pooled together to ensure that the number of cells was enough for downstream analysis. Cells were first subjected to a short low speed spin in a benchtop microcentrifuge (∼10 seconds) to separate T cells from spheroids. Cells in the supernatant were further centrifuged at 350 *g* for 5 minutes at 4 °C to remove the PBS and snap-frozen in dry ice. Samples were stored at -80 °C until metabolite extraction.

### Metabolite extraction and mass spectrometry analysis

CD8^+^ T cell collection and metabolite extraction were performed as previously described[36]. Polar metabolite abundances were measured using gas chromatography-mass spectrometry as previously described[37] with an 8860A GC system (Agilent Technologies) combined with a 5977C GC/MSD system (Agilent Technologies). The relative quantification of metabolite abundances was obtained by calculating the area under the curve of the corresponding peaks normalized to the area under the curve of the internal standard (i.e. D-Norvaline). The data were subsequently normalized according to protein content. Metabolite identification was performed by comparing the retention time, MS and MS/MS data with an in-house library and metabolite standards.

### Statistical analysis

Statistical analyses were performed in GraphPad Prism 11. Two-tailed paired Student’s t test was used for comparison of two paired groups. Ordinary one-way ANOVA test was used for comparison of multiple groups. Error bars display standard deviation (SD) unless otherwise stated.

## FIGURE LEGENDS

**Supplementary Figure 1: Recovery efficiency**.

CD8^+^ T cells embedded in PIC were recovered by liquefaction with cold PBS. The recovery efficiency was calculated using T cells in suspension as control. n=3 biological replicates.

