## Supplementary figures and images for "A thermoreversible hydrogel model for *in situ* and *ex situ* analysis of tumor-immune interactions"

### Supplemental Figure 1

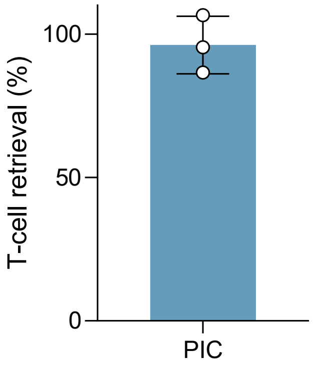

T-cell retrieval (%)
100
50
0
PIC
